# Nanowires unravel a time-correlated stochastic vectorial process in cells

**DOI:** 10.1101/2020.12.24.424270

**Authors:** Vishnu Nair, Matthew Seebald

## Abstract

A cell uses its cytoskeletal machinery to control its membrane projections to seek and obtain cargo from its microenvironment. Though this process has been studied extensively using spherical cargo, it remains largely unknown how the process operates with vectorial ones, which are non-spheroid rigid objects with an aspect ratio. In this study, a vectorial cargo, silicon nanowire, was observed to have multiple modes of initial contact and to realign along a membrane projection or on a lamella. Using a qualitative theoretical approach, we demonstrate how membrane energy fluctuations potentially drive this realignment of a vectorial cargo. This was understood by calculations which establish how aspect ratio controls the energy landscape in a vectorial object and its influence on relative energy stability of nanowire-membrane contacts. A study of the realignment transport of vectorial cargoes and their comparison with Ornstein-Uhlenbeck process simulations revealed how one-dimensional time-correlated noise manifested in the transport process. Furthermore, a comparison between sliding of nanowires on cell membrane contacts versus rotational realignment with the same model revealed identical characteristics behind both. The understanding that one-dimensional time-correlated noise underlies both sliding and rotation of a vectorial cargo establishes how cytoskeletal dynamics effectively couples their realignment with subsequent transport for phagocytosis. This work establishes the significance of vectorial cargoes and the nature of underlying vectorial processes that enable their cellular processing.

## Introduction

Cells explore their microenvironment by manipulating their membrane stochastically.^1,2^ This manipulation is done through the underlying cytoskeletal machinery which enables environmental exploration and is often triggered by cues received through the extracellular matrix (Figure 1a).^3-6^ These cues are stochastic forces generated by nearby or distant cargoes in the cell’s microenvironment.^3,7,8^ In response to these cues the cell uses its membrane to seek out these cargoes for various purposes (Figure 1b).^9-11^ Depending on the cell type and its physiological function, the fate of cargo a cell captures from its microenvironment varies.^12-15^ For example, a macrophage would try to digest bacteria in its lysosome after its phagocytosis.^16^ Though phagocytosis is generally well understood^17^, the transport processes involved in the capture of these cargoes by the cell membrane still remains an active topic of investigation.

**Figure 1.**
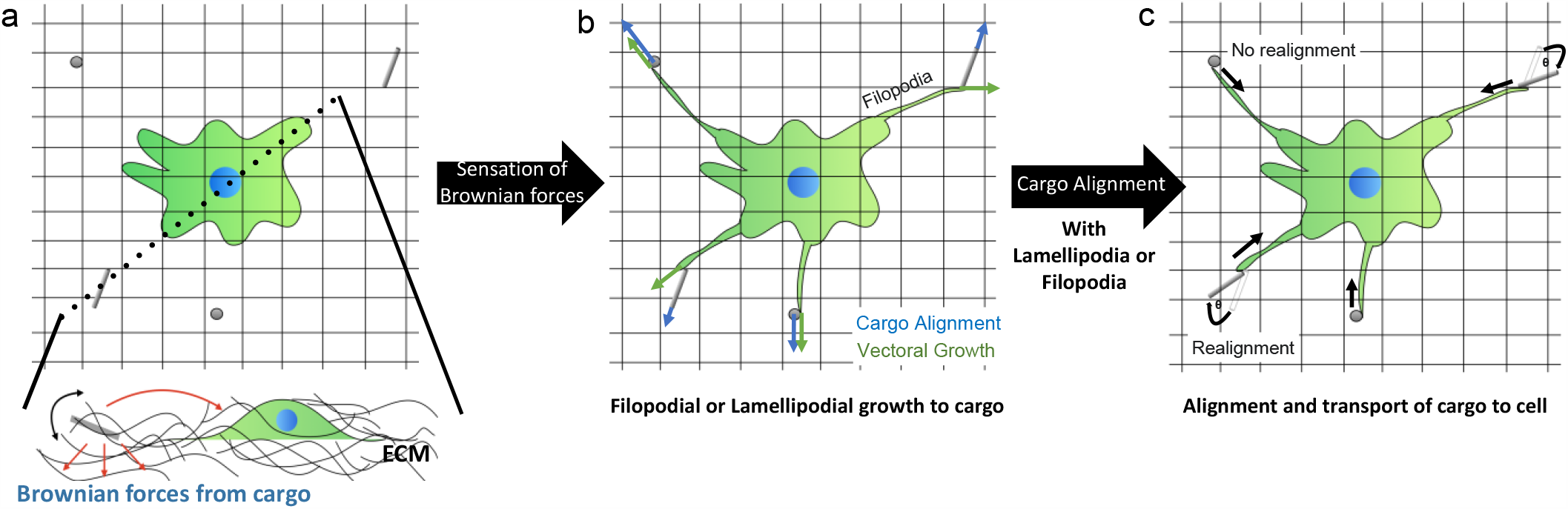
Hypothesis underlying this study: Cells are known to sense Brownian forces generated by objects lying in their microenvironments. Such a sensation is realized through the extracellular matrix (ECM), which transfers these forces to a cell (a). In response to the forces received from the ECM, a cell explores its microenvironment using membrane protrusions like lamellipodia or filopodia. The exploration is performed randomly until the membrane protrusion makes contact with the object (b). Once a contact is established between the membrane protrusion and the object, the cell transports it to the body for phagocytosis (c). However, the geometry of an object and its alignment with respect to the membrane protrusion should be considered. Unlike a sphere, which has only two translational degrees of freedom in plane, an object like a nanowire has an additional rotational degree of freedom. The nanowire and its initial alignment with membrane projection are a constraint for the cell when it comes to optimal membrane contact and transport, in comparison with a sphere. Thus, a cell realizes nanowire as a vectorial cargo and realigns it for optimal contact and subsequent transport (c).

Previous studies on inert nano or microparticle transport by filopodia and lamellipodia have been studied using optical traps.^18-21^ In studies using an optical trap, the particle of interest was placed on a filopodia or lamellipodia of a cell and the forces exerted on the particle studied by interferometric tracking. ^18-21^ However, the cell’s machinery for self-locating and transporting the particle weren’t utilized in these studies. Studies with particles have also revealed that contact made by membrane projections with the nano or microparticle often occur through a wrapping mode -where the projection wraps around the particle before retracting it towards the cell body- or in a lever arm retraction-where the projection pushes in the particle into the cell body.^19^ Besides these contact modes wherein the membrane projections have to move or retract themselves, a sliding motion of the particle without retraction of the membrane projections can also occur.^18^

Though spherical particles have been widely studied, it remains largely unknown how vectorial cargoes would behave in a similar context. By the term vectorial cargo, we refer to objects with an aspect ratio, for example a nanowire. Objects with an aspect ratio often make a larger contact with a cell membrane, compared to a sphere with an aspect ratio of one. Thus, intracellular factors which detect membrane curvature and stabilize variations in it should be considered.^22^ BAR proteins, which dimerize to stabilize curvature in membranes, have been realized to contribute significantly in this scenario.^22,23^ Moreover, BAR proteins are known to stabilize cylindrical curvatures better than spherical ones.^24,25^ This motivates one to think how a nanowire cargo would preferably be aligned along its cylindrical interface with the cell membrane. Hence this would place a demand that a nanowire would be realigned to such an interfacial geometry by the cellular machinery (Figure 1c).

The goal of this work is to test the nanowire realignment hypothesis arising from mechanotransduction that nanowire cargoes may cause. The dynamics, nature of the mechanical process, and the cellular machinery underlying this realignment process is widely unknown and hence of exploratory interest. In order to test this hypothesis, we picked VLS grown silicon nanowires as our choice of vectorial cargo. This is because silicon nanowires have previously been used to understand phagocytosis, rotations, and bending in various cells.^26-29^ Furthermore, silicon nanowires are biocompatible and biodegradable.^30,31^ Building on this hypothesis, we demonstrate how nanowires can cause primary fibroblast cells to perceive them in their environment, and form contacts to interface with them. Further, we observe and analyze realignments of nanowire cargoes by cells and establish, using fundamental statistical mechanics, how time-correlated noise underlies this phenomenon.

## Materials and Methods

### Cell culture

Primary rat cardiac fibroblasts were cultured from P1-P5 Sprague-Dawley Rats. The primary cell culture was carried out according to the manufacturer guidelines on the Pierce™ Primary Cardiomyocyte Isolation kit (Catalog No: 88281). The fibroblasts were separated from digested rat hearts by seeding them on plastic flasks, leaving cardiomyocytes in the supernatant. After an hour of seeding, the cardiomyocyte rich supernatant was removed, allowing fibroblasts to proliferate in the dish. The fibroblast cells were cultured in complete media consisting of DMEM with 5% Fetal Bovine Serum, 1% Pencillin-Streptomycin, and 1% Glutamax. The primary cell culture experiments were performed. The cell culture from rats were performed with approval from the University of Chicago Institutional Animal Care and Use Committee.

### Silicon nanowire synthesis

Silicon nanowires were grown using the Vapor-Liquid-Solid (VLS) growth method on silicon wafer chips. Citrate stabilized gold nanoparticles of 100 nm diameter were used for seeding the nanowire growth, which was performed at 470 °C and 40 torr for 15 minutes. The source of silicon in this CVD reaction was silane gas diluted with hydrogen carrier gas. The flow rates of silane and hydrogen were 2 sccm and 60 sccm, respectively. The nanowires were transferred from wafer chips to the cell culture media by ultrasonication.

### Live Imaging

Cells were seeded at a confluence of 10-15% on glass-bottom dishes (CellVis-D35-10-1.5-N) coated with Type 1 collagen (Corning). These cells were then transfected with CellLight™ Actin-GFP, BacMam 2.0 Reagent (Catalog Number: C10582) following the manufacturer’s protocol. Post transfection, the cells were treated with full media containing silicon nanowires. Live imaging was performed after nanowire treatment, in complete media made in Fluobrite DMEM, on a LEICA-SP5 STED CW confocal imaging system, and enclosed in an incubator at 37 C with 5% carbon dioxide. Actin fluorescence was recorded using a hybrid CCD detector. Nanowires were imaged by recording DIC images or reflected light, which was separated from the fluorescence using an acoustic optical beam splitter. Imaging was done at frequencies ranging from ∼0.1Hz to 1Hz, for a total duration of ∼3 to 10 minutes. The imaging frequency and total duration of imaging was limited by fluorescence bleaching occurring in that particular imaging frame.

### Fixed Imaging

Cells treated with nanowires were fixed, permeabilized and blocked using Image-iT™ Fixation Kit (Catalog No: R37602) following the manufacturer’s protocol. The cells were stained using ActinGreen™ 488 ReadyProbes™ Reagent (AlexaFluor™ 488 phalloidin) (Catalog No: R37110) following the manufacturer’s protocol. Actin fluorescence was recorded using a hybrid CCD detector. The nanowires were imaged by recording reflected light which was separated from the fluorescence using an acoustic optical beam splitter.

### Electron Microscopy

Nanowire treated cells were trypsinized and re-suspended in fresh media. These cell suspensions were then frozen at high pressure (BalTec) and embedded in HM20 Resin (Electron Microscopy Services) after freeze substitution (Leica AFS) in acetone along with 2% Uranyl acetate (Electron Microscopy Services). Resin embedded cells were ultramicrotomed (Leica) into 90 nm sections. The imaging was done on a Technai F30 Electron microscope at an accelerating voltage of 300 kV.

### Model for membrane energy calculations

Initially, when the nanowire makes contact with the membrane, its spherical cap region forms an interface with the filopodial membrane which bends to accommodate that spherical shape. Similarly, when the wire finishes its rotation and it becomes aligned with the filopodia, it lays flat on the membrane which bends to accommodate its lateral cylindrical surface. The nanowire can be considered to have smooth hemispherical ends as these are wires grown by VLS method.^32,33^

The overall changes in reduced (dimensionless) energy for a sphere and cylinder coming into contact with a membrane which subsequently deforms to their shapes were used to calculate the change in energy associated from a spherical to cylindrical topographical transformation as the wire rotates.^34-36^ These calculations are based on Helfrich theory. ^34-36^ The total reduced energy change (with respect to free standing planar membrane) can be expressed as a sum in the changes of reduced free energy, contact energy, and the energy of adhesion:

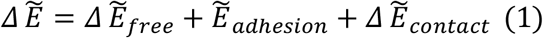

where, in the case of a cylinder 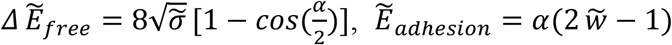 where *α* is the wrapping angle, 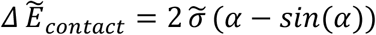 is the reduced adhesion energy per unit area and 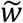 is the reduced membrane tension. For the cylinder, the reduced variables here are related to actual according to the following relationship: 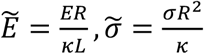 and 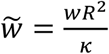, wherein κ is the bending constant of the membrane, R is the radius of the nanowire and L is the length of nanowire. The net energy change for a cylinder is hence:

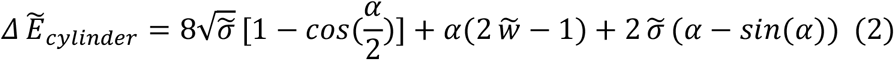

Similarly, for spheres 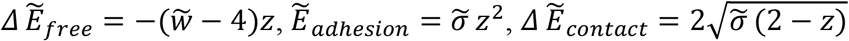 where z, in this case, is degree of wrapping: z = 1 − *cos*(*α*)

For the cylinder, the reduced variables here are related to actual according to the following relationship: 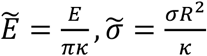 and 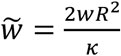, wherein κ is the bending constant of the membrane, R is the radius of the nanowire or sphere. The net energy change for a sphere is hence:

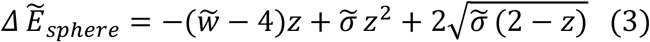

In order to understand the realignment of the nanowire, the overall change in energy following a spherical end-cap to cylinder transition via rotation was calculated by taking the difference between these two sums 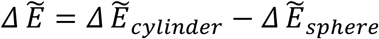. This difference 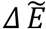 was plotted as a function of *α* is the wrapping angle and [inneq] is the reduced adhesion energy for three order of reduced tension 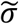.

Furthermore, in order to introduce aspect ratio dependance into our simulations we use the reduced energy expression 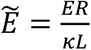 for cylinder and 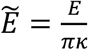 for sphere, to obtain E. In order to keep all variables in the dimensionless framework we use Boltzmann probability, considering the system as a two-state system (initial -spherical interface versus final-cylindrical interface)

Probabilities of being in each state were calculated using the Boltzmann probability function as:

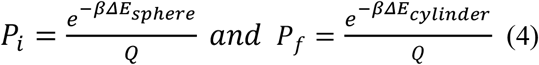

Where Q is the partition function given by 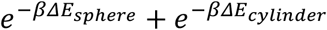. The values used for simulation were: 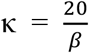 and aspect ratio 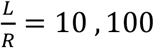, 100, *and* 1000, where R is in order of nanometers and L in the order of microns.^34^

### Calculation of dynamic variables for rotating wires

Rotation of straight, non-kinked silicon nanowires on a filopodia or lamella were analyzed. Such nanowires were identified and tracked using ImageJ. The axes of rotation were approximately identified by inspection and coordinates of either ends along with the center of mass obtained for each timestep. Schematic S1 explains a typical nanowire under analysis.

The information from tracking was used to determine the angle between the nanowire and outward projection of filopodia (Figure 1c) or the angle change with respect to initial configuration on a lamella at every timestep. This enabled generation of an angle of rotation as a function of time (i is the nanowire label or index): *θ*_*i*_(*t*)

Clockwise rotation was given a positive sign and anticlockwise rotation a negative sign. Angular velocity and angular acceleration of the wire were obtained by taking the first and second derivatives, respectively, of this angle function: 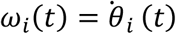 and 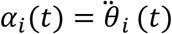

### Calculation of dynamic variables for sliding wires

To analyze linear sliding motion, the wire’s center of mass in the direction of motion was tracked at each timestep to construct a displacement function: *ri*(*t*)

The first and second derivatives were taken to find the velocity and acceleration, respectively:

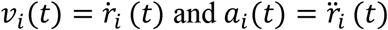

### Brownian analysis of rotating and sliding nanowires

Correlation time (*τc*) can be obtained by fitting an exponential decay function to the auto-correlation function of the noise in angular velocity^18^:

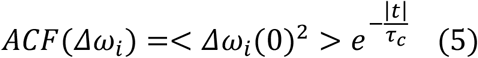

The noise in angular velocity was estimated by subtracting the time averaged angular velocity value from the angular velocity function. where: *Δω* _*i*_ = (*ωi*−< *ω* _*i*_ >)

In the case of linear sliding motion, a similar calculation was done using velocity^18^

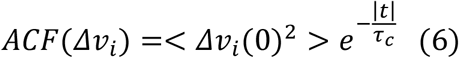

and *Δυ* _*i*_ = (*υi*−< *υ* _*i*_ >). It is to be noted that all measured angular velocity and linear velocity functions had a stochastic behavior but non-zero average for every nanowire observed. This suggested the presence of time-correlated noise or colored noise.

Further to estimate the effective diffusion coefficient of the nanowires which were tracked for rotations or sliding we used the following relationship for rotations and sliding,^37^ respectively:

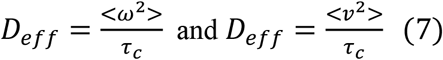

Finally, diffusivity (*α*_*i*_) of a given wire was extracted by fitting the following exponential relation to the rolling frame mean-squared displacement for rotations and sliding^29^, respectively:

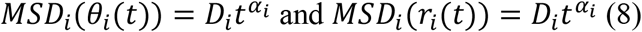

### Simulation of time-correlated stochastic process

To understand the nature of the noise correlation which underlies the observed rotational alignment and sliding transport of the nanowires, we use a model for planar motion of active point particles developed by Weber et al. and Radtke et al.^38-40^ This model takes into account the stochastic nature of surroundings such that the angle dynamics are driven by time-correlated noise, resulting in an effective diffusion coefficient. From this work, we specifically used the solutions to equations of motion involving systems with non-zero average angular velocity. According to this model, the angular velocity of a nanowire is subject to stochastic environmental noise, enabling the use of an Ornstein-Uhlenbeck process (OUP). In our system, the interface between the cell membrane with the wire served as the noisy environment, where the stochastic forces are transferred from cell to the nanowire. From this model, the angular dynamics for in plane motion could be written as a sum of angular velocity and noise:

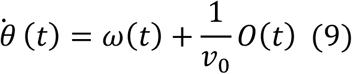

where *0*(*t*) represents the continuous-valued colored OUP noise, and v_0_ is the contribution to velocity of the center of mass from rotation. This contribution is given by: *υ* _0_ (*t*) = *x ω* (*t*)

In the case where the magnitude of measured value of *V*_*cm*_ (Net velocity of center of mass) and calculated value of *V*_0_ were within the same order, the motion of the wire was concluded to be mostly rotational. In the case of pure rotation *V*_*cm*_ and *V*_0_ are interchangeable and hence measured experimental *V*_*cm*_ was used for simulations.

A continuous-valued colored OUP noise is defined by the following relation:

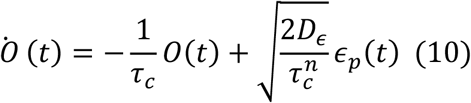

Where 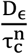 is the noise intensity, n is the dimension of D*ϵ* and n = 0, 1, or 2. The noise dimension n=0,1,2 correspond to processes in one, two and three dimensions in three-dimensional space, respectively. Gaussian white noise is given by *ϵ* _*p*_ (*t*) and *τ* _*c*_ is the noise correlation time. According to the Einstein Relation: *D*_ϵ_ = γ*k*_*b*_*T*, ^40^ where *γ* is the friction constant at cell -nanowire interface where motion happens,^18^ *k*_*B*_ is the Boltzmann constant, and *T* is the temperature. This equation enables the noise intensity to be related the heat bath (cell) which sources the noise. For < *ω* > ≠ 0 and trajectories (*θ* _*i*_ (*t*)) that satisfy the above system of differential equations (eq.9 and eq.10):

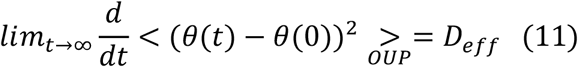

Where *D*_*eff*_ is the effective diffusion constant that arises out of this motion, as described by this model. The solution arrived at by Weber et al.^39^ in the limit of small correlation times 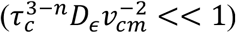 is:

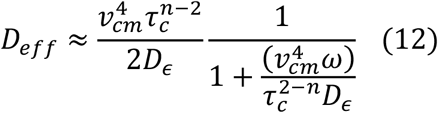

and in limit of large correlation times 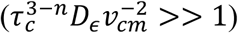 is

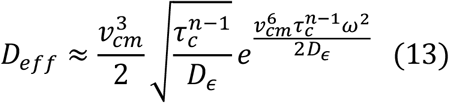

The simulations were carried out by an appropriate choice of values based on experimentally estimated dynamic variables. The experimental estimates are listed in Table S1 for rotating wires and in Table S2 for sliding wires. The simulation parameter values which were further obtained from literature and experimental estimates (Table S1 and S2) are listed in Table S3. For studies on rotating nanowires, the main variable is the angular velocity whose average value across wires (Listed as “Avg” in Table S3) along with the lower and upper limit (“L” and “U” in Table 3, respectively) are based on data in Table S1. The average velocity of center of mass was also obtained across wires from Table S1. In case of sliding wires, the angular velocity is zero and the main variable is the velocity of the center of mass. Similar to the rotational case the average value of velocity in the case of sliding wires (listed in Table S3) along with their upper and lower limit are based on Table S2.

Before we proceeded to compare the value of effective diffusion coefficient as a function of correlation time with experimental data, we used the simulation parameters in Table S3 to decide whether our problem lies in the limit of large or small correlation times. The inequality: 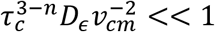 was tested using values from Table S3. For n=0, the inequality gives *τ* << 10^7.5^. For n=1, the inequality gives *τ* _*c*_ << 10^8.5^. For n=2, the inequality gives *τ* _*c*_ << 10^15.5^. This test suggests that the correlation time required for large correlations based on parameters is extremely high. Hence our problem completely lies in the domain of small correlation times and the expression valid for our problem would be equation 12.

This expression was simulated for n=0, 1, and 2 under parameters in Table 3 with variables as angular velocity in the case of rotations and linear velocity in the case of sliding. The simulations were plotted on a log_10_-log_10_ scale, with log_10_(D_eff_) on y axis and log_10_(*τ*_*c*_) on x axis. The experimentally obtained effective diffusion coefficient and correlation times (As explained in Brownian analysis section of Materials and Methods) are plotted as points superposed on the simulation plots.

## Results and Discussion

### Formation of cell-nanowire contacts and phagocytosis

We first cultured fibroblast cells at low confluence. Fibroblasts were chosen cell type as they are well understood and widely used in the field of mechanobiology and extra-cellular matrix biology.^41-43^ These low confluent cells were then seeded with nanowire in full media and incubated until various time points. Live imaging is started within an hour of seeding as soon as the nanowires settle down. At this instance, we observed the formation of lamellipodia in the direction of the nanowires near vicinity of the cell (Figure 2a). Similarly, filopodia also initiated contacts with nanowires (Figure 4a). Fixed cell imaging of cells, 12 hours after seeding of the nanowire revealed unique geometries of filopodia -nanowire contact (Figure 2b-e). Filopodia were observed to be wrapping around a nanowire on its edge (Figure 2b, d) or holding wire using multiple projections (Figure 2c, e). Alongside these, we also observed, using electron microscopy (after 12 hours of seeding nanowire), that lamellipodia pushed nanowires into plasma membranes for uptake (Figure 2f). The wrap-around and push-in modes of contact have been previously observed with spherical nanoparticles, however the holding mode is unique to a nanowire.^19^ This is potentially due to the fact that multiple projections are required to physically hold an elongated object. These results confirm the idea that cells are responsive towards nanowires in their environment and adopt geometrically sustainable contact modes to interact with them.

**Figure 2.**
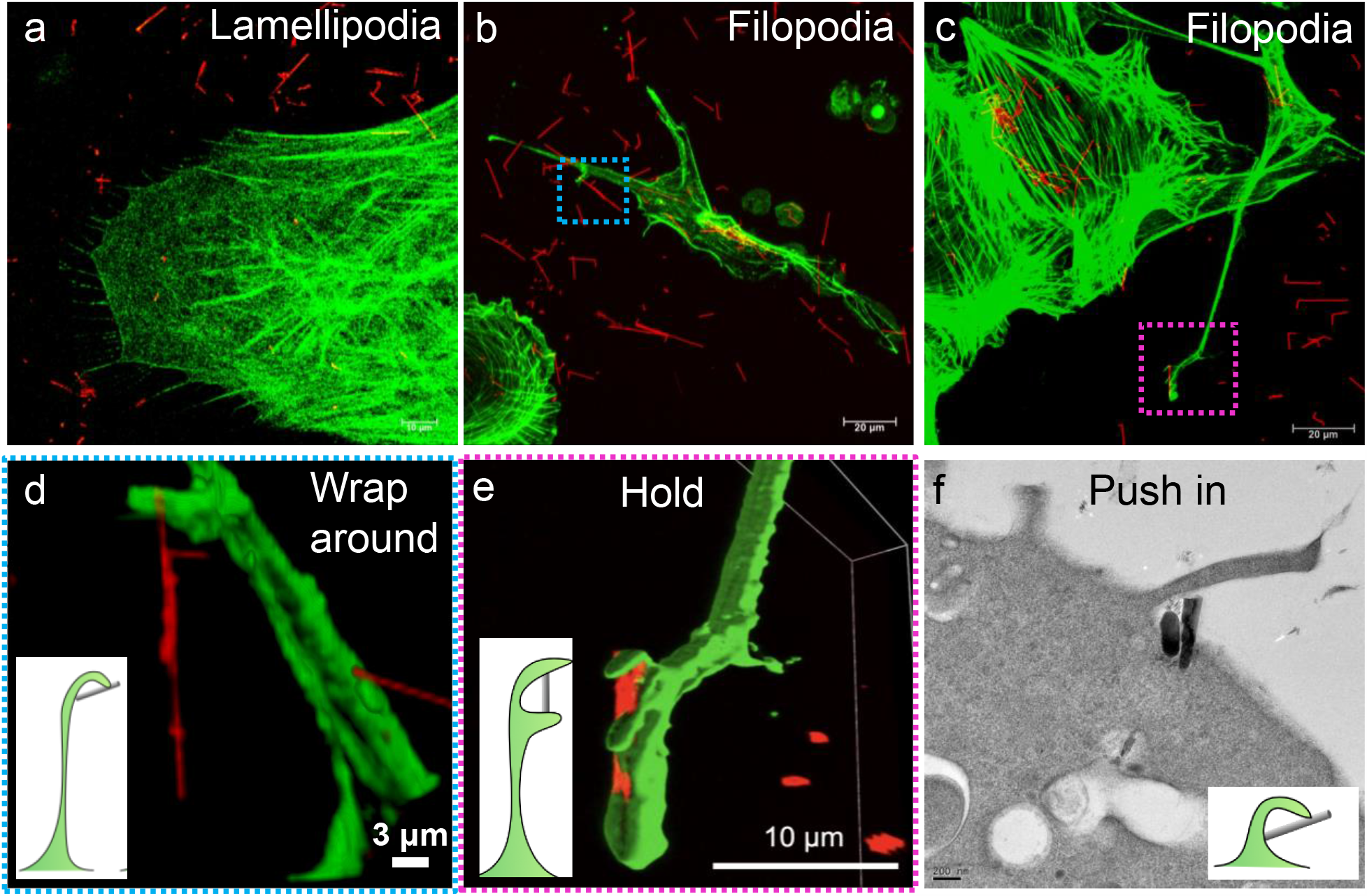
Formation of cell-nanowire contacts through lamellipodia and filopodia: a. Live confocal image taken an hour after seeding nanowires. The image shows a cell using lamellipodia to contact various nanowires in its vicinity. b and c. Fixed confocal image showing filopodia-nanowire contacts, 12 hours after nanowire seeding. d. Enlarged 3D reconstruction of boxed area in b showing a membrane wrap around configuration of contact between cell and nanowire. e. Enlarged 3D reconstruction of boxed area in c showing membrane projections from a filopodia holding onto a nanowire. f. TEM image showing a membrane projection pushing a nanowire into the plasma membrane.

**Figure 3.**
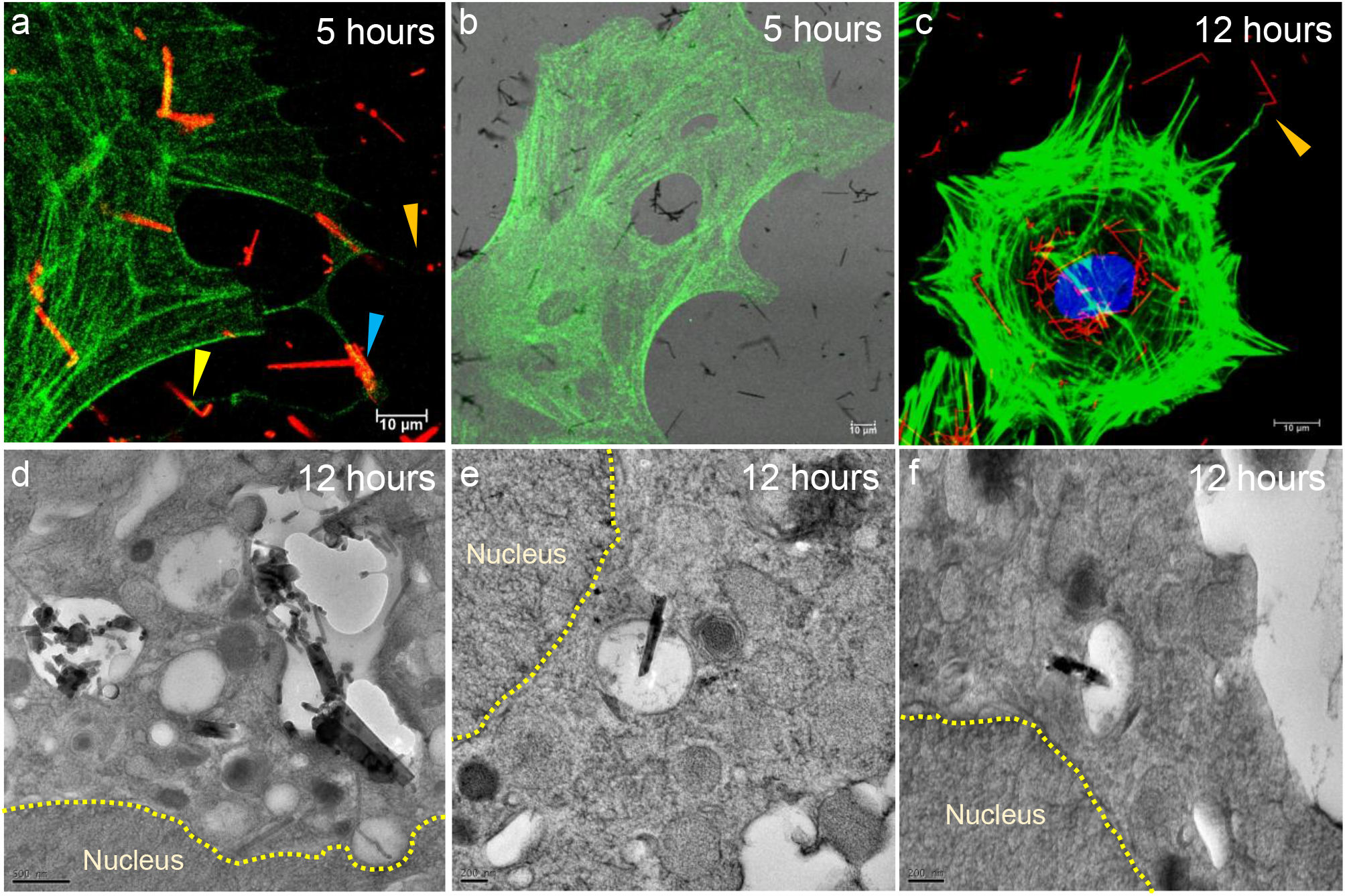
Fate of nanowires: Live cell imaging showing nanowire phagocytosis. The nanowire is viewed in reflection channel (a) and DIC channel (b). (c). Fixed cell confocal image showing continuous uptake and long-term phagocytosis leading to nanowire accumulation around the nuclear region. Long term nanowire phagocytosis was studied using electron microscopy showing nanowires in vesicles located near the nucleus-(d) and (e). (f). Titled view of the section in (d) showing that the nanowire is internalized in the cell.

**Figure 4.**
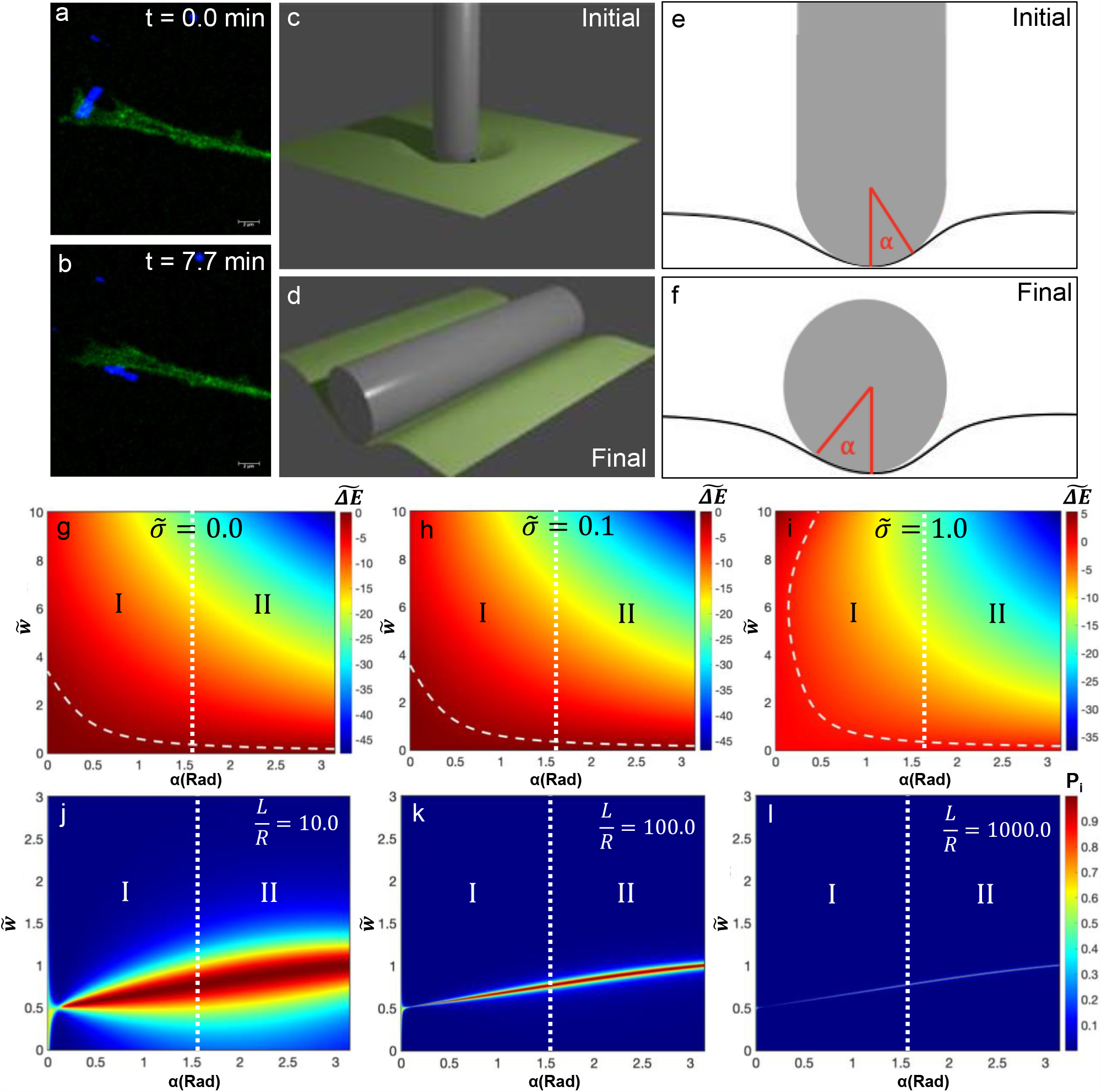
Preliminary observation and simulation supporting the nanowire realignment hypothesis: A nanowire initially aligned an angle to a filopodia (a), eventually aligns parallel to its surface (b). The initial (a) and final configuration (b) of a nanowire on a membrane can be modelled as in (c) and (d), with cross-sections as in (e) and (f) respectively. Since the nanowires are CVD grown, the edges of nanowire are approximately hemispherical as illustrated in (e). Hence the initial mode of contact can be modelled using a spherical surface whereas the final, using the lateral surface of a cylinder. (g-i). Dimensionless energy difference 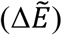 between final (d,f) and initial (c,e) configurations as a function of adhesion energy-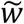 and wrapping angle-α with increasing tension -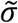. The dashed curved line indicates the 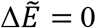 contour and zone I the pre-phagocytotic and zone II the phagocytotic regime of wrapping. The 2D heatmaps of 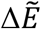 (g-i) show how significant tension 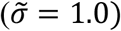 is necessary to obtain a huge range of energetically feasible configurations required for the observed realignment. The Boltzmann probability of initial configuration as a function of adhesion energy-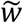 and wrapping angle-α with a tension - 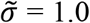with an increasing aspect ratio from 10 (j), to 100 (k) and finally 1000 (l). The probability distributions show how an increasing aspect ratio (L/R) decreases the number of allowed configurations for spherical contact, thus favoring realignment to a cylindrical contact.

The observations in Figure 1 suggest the responsiveness of the cells towards nanoscale objects in their environment. However, five hours post seeding the nanowire, live imaging revealed nanowires in vesicles (Figure 3a, 3b). Based upon previous reports in literature this observation is due to phagocytosis of the nanowire.^26,27^ Even during the process of phagocytosis, we observed lamellipodia making contact with nanowires and filopodia with aligned (Figure 3a -Orange and Blue arrow) or entangled nanowires (Figure 3a-yellow arrow) being transported towards the vesicle. Similar observation was made in fixed cell imaging with cells incubated with nanowires for 12 hours (Figure 3c). After twelve hours, lamellipodia were making contact with nanowires (Figure 3c – orange arrow) though many potentially internalized nanowires were clustered around the nucleus. Electron microscopy performed with cells incubated with nanowires for 12 hours confirmed these results, as nanowires internalized in vesicles were found around the nucleus in large quantities (Figure 3d,3e). Furthermore, tilting experiments revealed a three dimensional view of the vesicle with a nanowire inside them (Figure 3 e,f). These observations confirm the fact that the final goal of cells sensing, initiating contact, and transporting nanowires is for internalization by phagocytosis.

### Nanowire realignment on cell membrane

The hypothesis behind this study is that nanowires being a vectorial cargo, as opposed to nanoparticles, which are spheroid, would lead to cells realigning them to optimize their contact with wires for transport and phagocytosis. Thus, we first attempted live imaging to observe this phenomenon (Figure 4 a,b). In this specific case a nanowire making an angle with the filopodia is rotated to align parallel to it. Since filopodia and lamellipodia are both extensions of the cell membrane, we use a general theoretical approach to understand potential reasons behind this observed realignment. Considering a CVD grown nanowire to be nearly hemispherical at its ends, rather than sharp, we look at membrane bending due to nanowire contact. The initial configuration can be considered as a spherical interface with only the end of nanowire making contact (Figure 4 c, e). The final configuration can be a cylindrical interface with the lateral surface of the nanowire making contact (Figure 4 d, f). The variables of membrane energy are the wrapping angle (α), the membrane tension (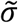) and the adhesion energy 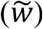, according to the model specified in materials and methods. We investigated the energy difference 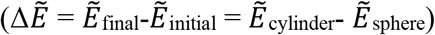 between initial and final configurations subject to these dimensionless variables. Additionally, we used the Boltzmann probability of the initial configuration (P_i_ = P_i_ 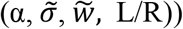 to understand the probable contact modes (A contact mode corresponds to a set of variables - α,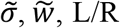) which allow the initial configuration.

The energy difference 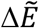 plotted as a heat map of adhesion energy and wrapping angle for various orders of tension (Figure 4g-i) show that most contact modes energetically favor 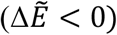 the cylindrical interface (final configuration) over the spherical interface (initial configuration). However, there is only a significant difference in the number of energetically favorable contact modes at 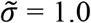 compared to 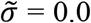 and 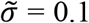. This difference is observed by looking at the 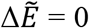 contours overlayed on the simulation. Since real cell membranes have significant tension that controls its dynamics,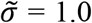 is a condition that could potentially serve as an experimental equivalent.^34,35^ These energy plots reveal more negative 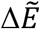 values and more contact modes (More area for 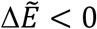) for wrapping angles more than 90° (Region II), wherein the membrane completely wraps the object for phagocytosis. This suggests two ideas-First, a cylindrical interface is favored for phagocytosis against a spherical interface, as the plot mainly has 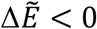 regions. Second, phagocytosis (Region II) has more energetically favorable modes than during contact or transport (Region I), which drives the ultimate phagocytotic fate of the nanowire. Thus, a realignment to a cylindrical interface is energetically favored for further transport and phagocytosis. However, generally, more negative 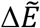 values for increasing variables (α, 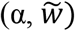) in the plot suggest that realignment could also be validated by the relative stability between any initial and final mode of contact such that 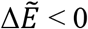.

Furthermore, to explore the role of aspect ratio in realignment, Boltzmann probabilities of the initial configuration are plotted as a heat map of adhesion energy and wrapping angle for various orders of aspect ratio (Figure 4j-l). These plots indicate that the allowed contact modes of a spherical interface decrease with increasing aspect ratio. This suggests that realignment is a prerequisite for a vectorial cargo to optimize contact interface prior to subsequent transport and phagocytosis. Besides introducing the requirement for realignment of vectorial cargoes, these simulations bring forth a general idea that vectorial transport like rotational realignment can happen generally at any nanowire-membrane interface. Furthermore, alignment of a nanowires entire length along a membrane may not be necessary to attain an energetically stable configuration.

### Rotational realignment at nanowire-membrane contacts

As understood using simulations, a vectorial cargo may require realignment to obtain an energetically stable interface with the membrane. Within the framework of this work, we were using nanowires which were expected to align to a cylindrical interface with the membrane. Hence this phenomenon should be observable on filopodia and lamella. Further screening of live imaging. enabled us to observe nanowire rotations happening on filopodia (Figure 5a-f) and lamella (Figure 5g-i, S1). We picked seven such rotating nanowires from three independent experiments for analysis to determine the nature of the process causing such rotations. The nanowires chosen for the study were linear and non-kinked to keep the analysis of their dynamics simple. The nanowires were then tracked for rotations with respect to their initial alignment or position. The angle of rotation, tracked with respect to initial alignment, can increase or decrease with time depending on the direction of rotation and corresponding sign convention (Figure 6a). The first-time derivative of the rotation with time gave us the angular velocity and the second time derivative, the angular acceleration. The angular velocity and the angular acceleration showed a stochastic behavior (Figure 6b, S2, S3) in their direction but possessed a non-zero average. This suggests that the forces involved in directing the rotation have a stochastic part along with a constant part. Such an active rotation in a specific direction, with underlying stochasticity, can potentially result from time-correlated noise.

**Figure 5.**
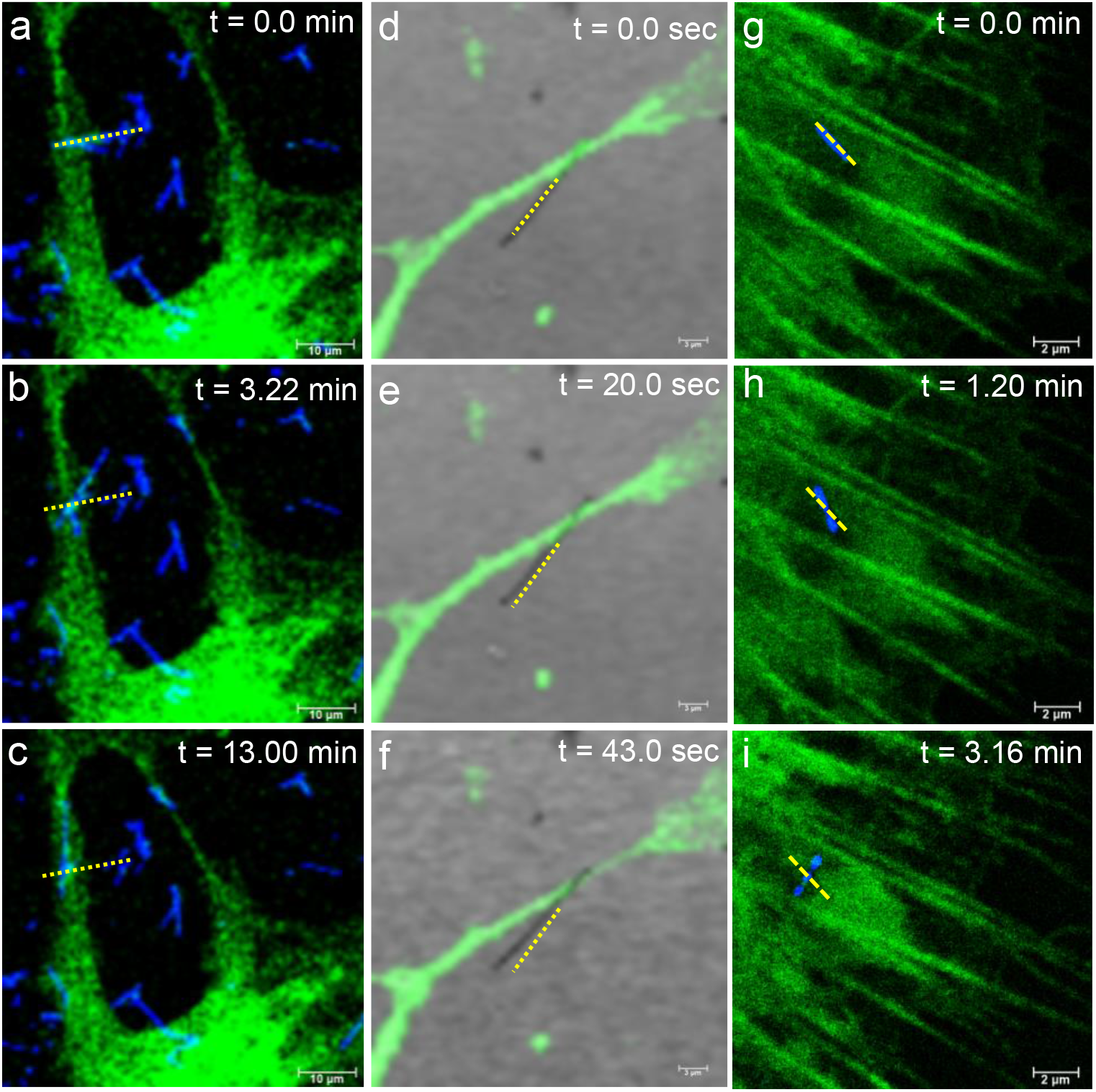
Representative data showing rotational realignment of nanowire: Snapshots from a nanowire aligning with a filopodia (a-c) and (d-f) and realignment on a lamella (g-i). The dashed yellow line represents the initial alignment of the nanowire.

**Figure 6.**
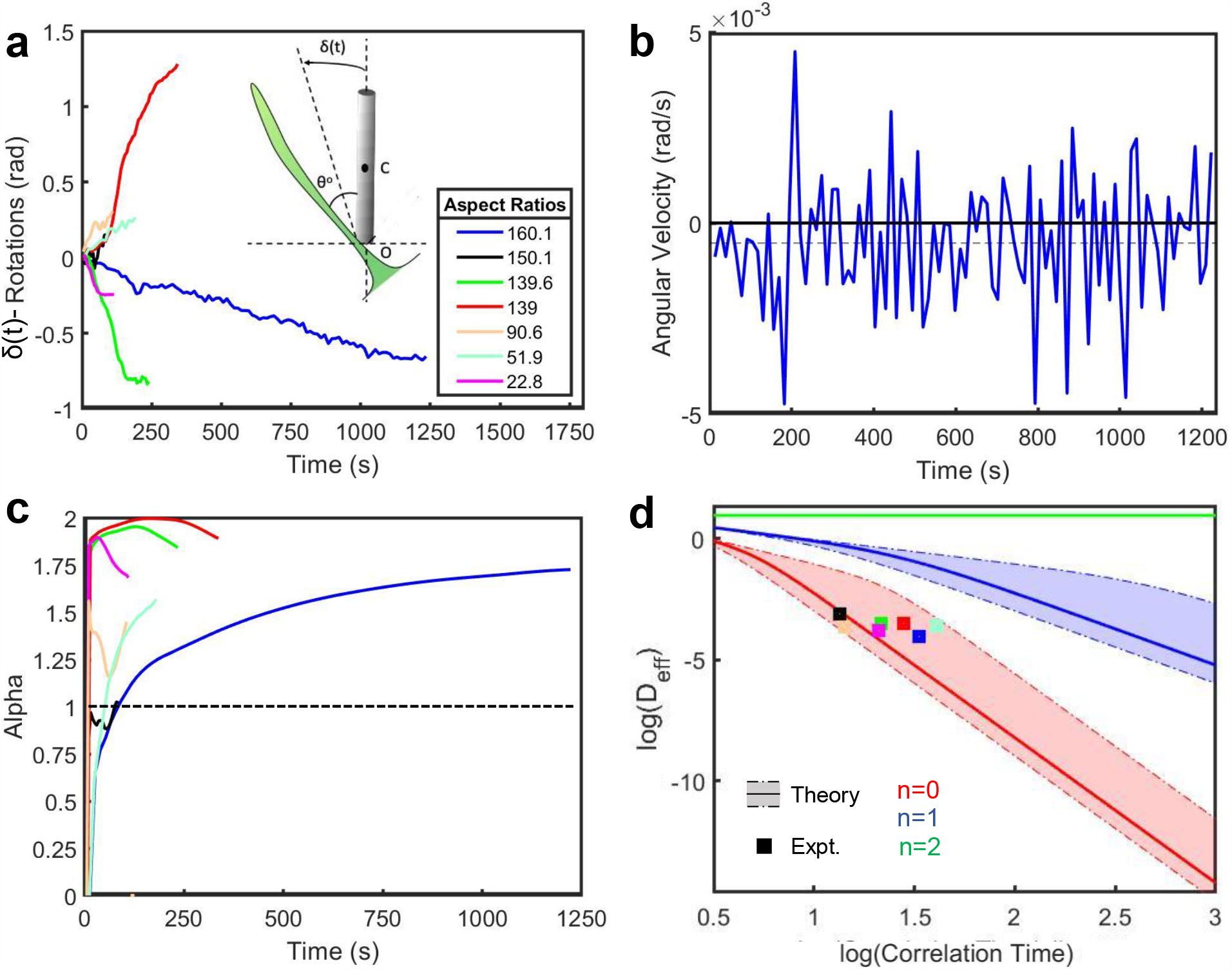
Analysis of rotating nanowires: a. Tracking measured rotation of nanowires with respect to their initial configurations as illustrated in the inset schematic. All traces in the figure are color coded with respect to their aspect ratio as in legend b. Representative first time derivative of rotations in (a) to obtain angular velocity of a rotating nanowire showing non-zero-time average (dashed line) alongside random flipping in direction. c. Anomalous exponent of various rotating nanowires showing an increasing super-diffusive with time suggesting an active transport performed by cell on the nanowire. Since the exponent value hadn’t saturated and fallen below one, it suggested that the nanowires being tracked were in the pre-phagocytotic regime. d. Comparison of effective diffusion coefficient of rotating nanowires with simulation of noise-correlated rotations with various dimensions of noise (n=0 corresponds to one dimensional noise, n=1 corresponds to two-dimensional noise and n=2 corresponds to three-dimensional noise.)

Analyzing these rotation processes using a Brownian diffusion model, the anomalous exponent as a function of time grew above one for all wires (Figure 6c). The growth of the anomalous exponent above one suggests the phase where cell membrane establishes a contact with the nanowire.^26,29^ During the course of tracking the anomalous exponent, it remains above one suggesting active (super-diffusive) motion.^26,29^ Furthermore, the anomalous exponent remaining above one suggests that phagocytosis hasn’t been initiated in any of these tracked wires. ^26,29^ This conclusion is due to the fact that during phagocytosis the anomalous exponent falls below one to the sub-diffusive regime, which is not observed within the time frame of consideration.^26,29^ Given that basic Brownian analysis of these nanowire rotations suggested an active motion and pre-phagocytotic transport, the exact nature of the motion needed to be understood. Based on the model described in the Materials and Methods, we compared the effective diffusion coefficient (D_eff_ (τ)) simulated under experimentally relevant range of variables to effective diffusion coefficient obtained from tracking experiments. In accordance with the model in use here the effective diffusion coefficient is considered as a function of the correlation time in noise. The comparison is performed by overlaying average experimental data points with the simulation for D_eff_ (τ) as in equation 12 with average value of variables (solid line in Figure 6d) and their ranges (dashed-dot line in Figure 6d).

The overlay shows that for wires of various aspect ratio the values from tracking experiment fell well within the order of simulated values (Figure 6d). This match confirms that the time-correlated noise model held true in our system and that it explains the rotational motion the cell performs on the nanowire. Furthermore, the match between simulation and experiment happened specifically for the model involving noise in one dimension. Hence, this experiment suggests that one-dimensional time correlated noise drives the rotational realignment of nanowires on a membrane to attain lowest energy of the nanowire-membrane interface.

### Comparison of rotational realignment with sliding of nanowires

From the analysis of rotational dynamics of nanowires in previous analysis, we understood that they are driven by one-dimensional time-correlated noise. There is a significant possibility that such a one-dimensional stochastic velocity is generated by dynamics of actin assembly.^5^ This is because the machinery a cell possesses to produce localized one-dimensional stochastic velocities lie in dynamic instability of actin.^44,45^ The correlated-stochasticity underlying these rotations could potentially arise from that the dynamic instability in actin due to its random polymerization - depolymerization cycles.^5,11,44,45^ Since it has been understood that linear translational forces have been generated by actin dynamics, the exact mechanism of the coupling of such forces at membrane interfaces are highly complex and unknown.^19,20^ However, it’s possible using our model to order to draw a parallelism between the machinery underlying rotational realignment and linear sliding motion on cell membrane. Towards this goal, we analyzed nanowires using the same methodology and model used in the case of rotating nanowires. Wires sliding along a lamella and filopodia were tracked (Figure 7 a-f). In either case a close observation of the membrane-nanowire contacts during live imaging revealed how actin rich projections modulate as wires translate. It was observed that the actin projections rear (with respect to the direction of translation) to the nanowire (Red arrow in Figure 7 a-f) retracted and expanded forward, pushing the nanowire in the direction of motion. This observation reveals how an actin network performs a spatiotemporally correlated series of one-dimensional depolymerization and subsequent polymerization purely in a direction perpendicular to the wire, to produce motion. Though the exact internal process is very complicated and beyond the scope of the work, qualitatively we can observe the process of translation arising from dynamic instability of actin. Furthermore, this establishes a rational for the comparison between rotational realignment and linear sliding motion to correlate the nature of underlying processes.

**Figure 7.**
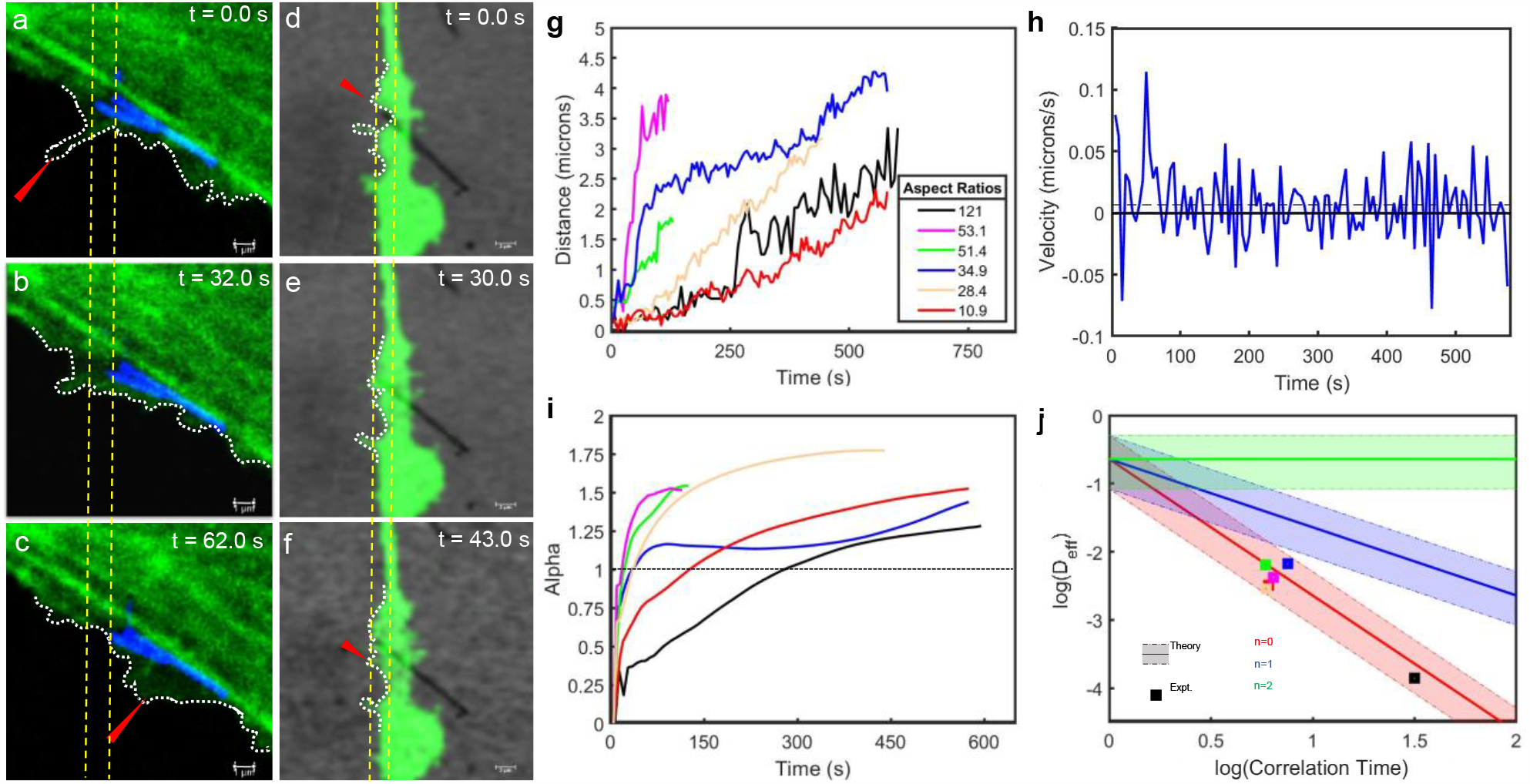
Representative data showing sliding of nanowire and their dynamic analysis: Snapshots from a nanowire sliding along a lamella (a-c). When the nanowire slides along the lamella a rear projection of the membrane (red arrow) that contacts the membrane retracts and expands forward causing the nanowire to move ahead. Snapshots from a nanowire sliding along a filopodia (d-f). A behavior similar to one for lamellar transport was observed here where in a rear projection retracted and expanded causing the nanowire to displace on the filopodia. g. Tracking measured displacement of nanowires with respect to their initial position. All traces in the figure are color coded with respect to their aspect ratio as in legend. h. Representative first time derivative of displacement in (g) to obtain velocity of a sliding nanowire showing non-zero-time average (dashed line) alongside random velocities. i. Anomalous exponent of various sliding nanowires showing an increasing super-diffusive with time suggesting an active transport performed by cell on the nanowire. Since the exponent value hadn’t saturated and fallen below one, it suggested that the nanowires being tracked were in the pre-phagocytotic regime. j. Comparison of effective diffusion coefficient sliding nanowires with simulation of noise-correlated translation with various dimensions of noise (n=0 corresponds to one dimensional noise, n=1 corresponds to two-dimensional noise and n=2 corresponds to three-dimensional noise)

Nanowires sliding along a filopodia and lamella (Figure 7a-f, S4) were tracked for displacements (Figure 7g). The first and second time-derivative of displacements gave the velocity (Figure 7h, S5) and acceleration (Figure S6). The velocity and acceleration both show a stochastic nature in their direction but with a non-zero average value. This characteristic was very similar to the observation in rotations. A basic Brownian analysis reveals that the anomalous exponent grows to one and saturates above one. This suggested that we were tracking wires in a time frame wherein the wires were attaching to the cell membrane (growth of alpha to 1).^26,29^ Besides, by not dropping to a value below one, the saturation suggested active (super-diffusive) transport before the start of phagocytosis (Figure 7i).^26,29^ Finally, we invoked the model used for simulating effective diffusion coefficient as in case of rotations. Simulations were performed within range of variables corresponding to the case of translating wires and compared with the results from tracking experiment. A comparison between simulation and experimental values of effective diffusion coefficient as a function of correlation time in noise revealed a significant overlap (Figure 7j) for n=0. Hence, we were able to conclude that even linear translations performed by cell membranes on nanowires were driven by time-correlated noise of one dimension. However, it is to be noted that the correlations times of noise have been observed to be an order longer in rotations than sliding motion (Table S1, S2, Figure S8). These observations enabled us to conclude that the rotational realignment and linear translations are driven by similar processes and potentially by the same machinery behind actin dynamics. Furthermore, this similarity underlying a vectorial and non-vectorial transport process suggests the connection between realignment of nanowire and their subsequent sliding transport required before phagocytosis.

### Conclusions and Future directions

Helfrich theory-based calculations demonstrate that relative energy stabilization could potentially drive rotational realignment of a nanowire at its interface with a cell. Besides this theoretical calculation, the phenomenon was observed through live imaging. Tracking the live motion and analyzing them revealed the process to be obeying a one-dimensional Ornstein-Uhlenbeck process. Furthermore, a comparison of the rotational realignment with sliding of wires revealed the underlying process to be similar. This potentially suggests that fundamental actin dynamics potentially enables contact, realignment and transport of the nanowires for phagocytosis. This work introduces a fundamental idea of how cells perceive vectorial cargo and the necessity of a vectorial process such as a rotational realignment for their subsequent cellular processing. Through this work we introduce the concept of a vectorial cargo and vectorial process in cellular biophysics. Though this work provides a simplified model of these concepts, it opens doors for future investigation for a deeper understanding of this phenomena. The membrane energy model could be made quantitive, if experimentalists in future estimate accurately the membrane tension, adhesion energy and developments in super resolution microscopy enable imaging of the nanoscale interface. The stochastic model used in this work treats rigid objects as particles for analysis. Thus, the dependance of dynamic variables like correlation time and diffusion coefficient on aspect ratio can’t be captured by these simulations. This brings forward the requirement of a theoretical framework to be built for Brownian dynamics of rotating rigid bodies. Hence this work opens an avenue for theoretical and experimental research on physics of biointerfaces.

## Acknowledgements

The authors acknowledge Dr. Bozhi Tian for his support and advice on this work. V.N. expresses his gratitude to Dr. Yamuna Krishnan for her feedback and suggestions in shaping this project. The authors acknowledge light microscopy core and advanced electron microscopy core facilities at the The University of Chicago, in this work.

## Author Contributions

V.N. designed the project, performed experiments and built the theoretical framework. M.S. performed all the simulations and theoretical calculations. V.N. and M.S. wrote the manuscript. V.N. and M.S. contributed equally to this work. The authors declare no competing financial interests.

