## Supplemental Information for "Nanowires unravel a time-correlated stochastic vectorial process in cells"

Vishnu Nair* and Matthew Seebald,
Department of Chemistry, The University of Chicago, Chicago, IL, USA

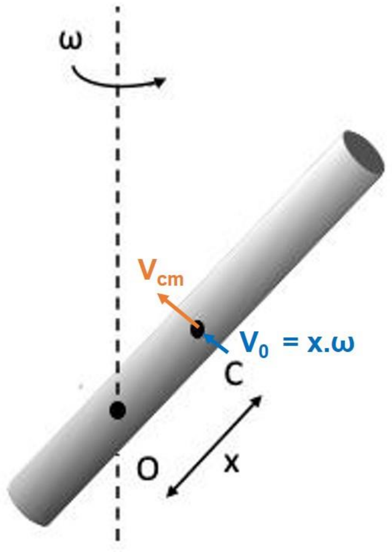

**Schematic S1**: Illustration of a nanowire with centre of mass C rotating along an axis passing through O. The axis of rotation is perpendicular to the plane in which the nanowire is rotating. The net measured velocity of the center of mass is $V_{cm}$ and the estimated contribution to it from rotational motion is $V_0$. x is the separation between the axis of rotation and the center of mass measured along the wire.

| Wire #- i | AR | Color | <ω> (rad/s) | <α> (rad/$s^2$) | <Vcm> (m/s) | x (μm) | Vo = xω (m/s) | $\tau_{corr}$ (sec) | D (rad$^2$ /s) |
|---|---|---|---|---|---|---|---|---|---|
| 1 | 139.60 | Green | -3.60E-03 | 1.71E-05 | 3.80E-08 | 6.98 | 5.08E-08 | 2.16E+01 | 3.13E-04 |
| 2 | 160.10 | Blue | -5.26E-04 | 2.26E-06 | 4.95E-09 | 8.05 | 8.21E-09 | 3.32E+01 | 9.40E-05 |
| 3 | 139.00 | Red | 3.80E-03 | -7.76E-06 | 4.07E-08 | 7.55 | 7.40E-09 | 2.79E+01 | 3.10E-04 |
| 4 | 95.40 | Pink | 1.30E-03 | -3.23E-05 | 6.07E-09 | 4.77 | 2.45E-09 | 2.39E+01 | 1.34E-03 |
| 5 | 150.10 | Black | 1.80E-03 | -2.04E-06 | 1.40E-08 | 7.50 | 2.75E-08 | 1.35E+01 | 8.24E-04 |
| 6 | 90.60 | Orange | 2.60E-03 | -3.10E-05 | 5.12E-08 | 4.53 | 2.32E-08 | 1.42E+01 | 2.23E-04 |
| 7 | 51.90 | Cyan | 1.40E-03 | -5.90E-06 | 4.77E-09 | 2.59 | 7.25E-08 | 4.07E+01 | 2.47E-04 |

**Table S1:** Average dynamic variables estimated from analysis of rotating wires

| Wire # - i | AR | Color | <$V_{cm}$> (m/s) | <a> (m/$s^2$) | $\tau_{corr}$ (sec) | D (μm$^2$/s) |
|---|---|---|---|---|---|---|
| 1 | 51.4 | Green | 1.15E-08 | -2.85E-04 | 5.88E+00 | 6.47E-03 |
| 2 | 34.9 | Blue | 6.70E-09 | -2.44E-04 | 7.52E+00 | 6.69E-03 |
| 3 | 10.9 | Red | 3.70E-09 | 1.36E-04 | 6.06E+00 | 3.26E-03 |
| 4 | 53.1 | Pink | 3.09E-08 | -6.25E-04 | 6.42E+00 | 4.24E-03 |
| 5 | 28.4 | Orange | 6.90E-09 | -3.49E-04 | 5.90E+00 | 2.85E-03 |
| 6 | 125.8 | Black | 5.20E-09 | 2.27E-05 | 3.17E+01 | 1.40E-04 |

**Table S2:** Average dynamic variables estimated from analysis of sliding wires

| Variable | ROTATION |
| --- | --- |
| $D_\varepsilon = \gamma K_b T$ | $4.55 \times 10^{-32.5}$ |
| $\gamma$ (Ns/m) | $10^{-7.5}$ (Ref.18) |
| $K_b$ (J/K) | $1.38 \times 10^{-23}$ |
| T (K) | 330 |
| $\omega$ (rad/sec) | Avg:0.0021<br>U: 0.005<br>L: 0.0001 |
| Vcm (m/sec) | $2.20 \times 10^{-08}$ |
| **Variable** | **SLIDING** |
| $D_\varepsilon = \gamma K_b T$ | $4.55 \times 10^{-32.5}$ |
| $\gamma$ (Ns/m) | $10^{-7.5}$ (Ref.18) |
| $K_b$ (J/K) | $1.38 \times 10^{-23}$ |
| T (K) | 330 |
| $\omega$ (rad/sec) | 0 |
| Vcm (m/sec) | Avg:$1.08 \times 10^{-08}$<br>U: $1.25 \times 10^{-09}$<br>L: $0.75 \times 10^{-08}$ |

**Table S3:** Experiment derived parameters for simulation of noise-correlated stochastic process.

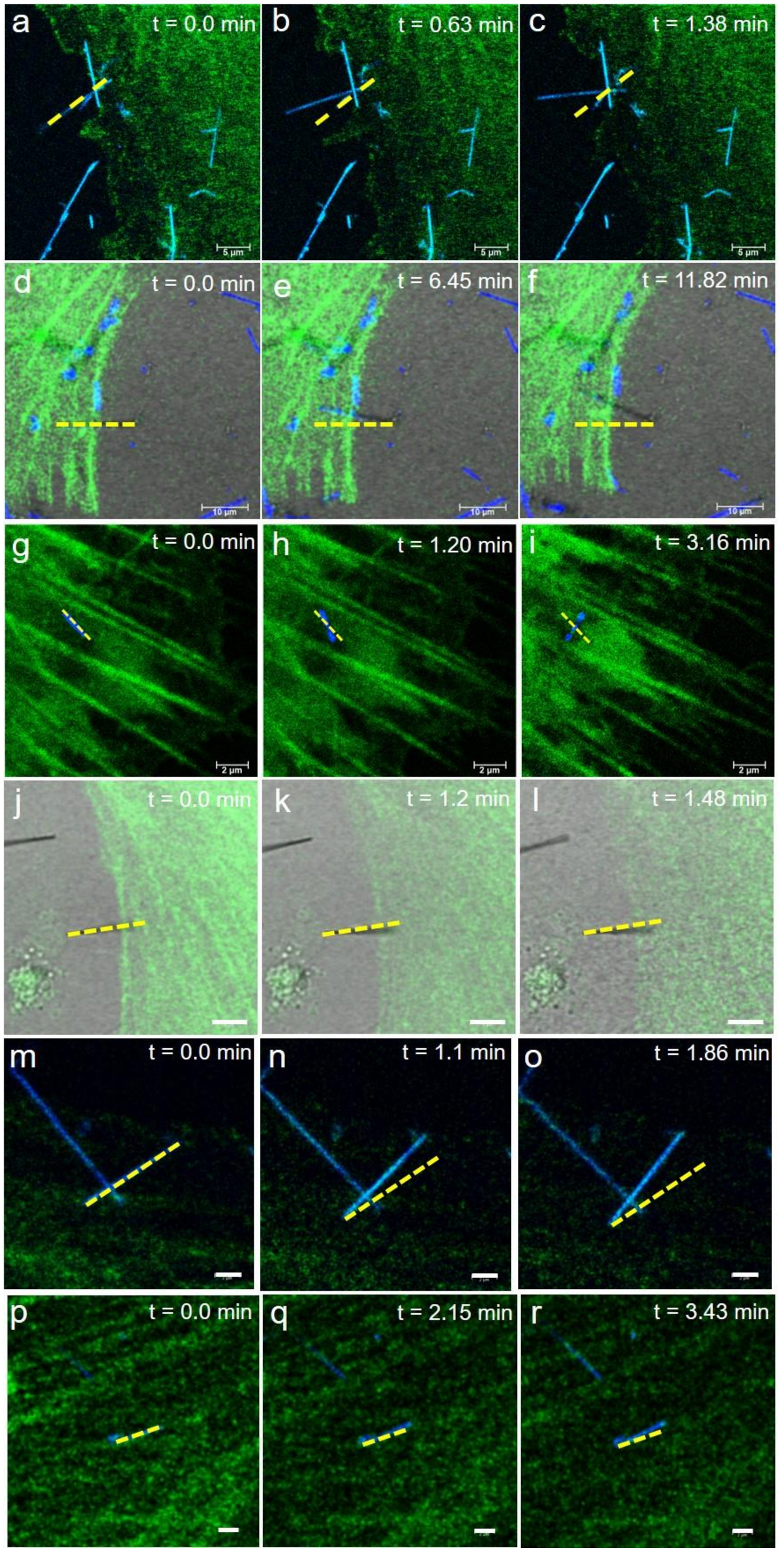

**Figure S1:** Snapshots of nanowire rotating on a lamella. Nanowires correspond to respective # as listed in Table S1: (a-c) -#1, (d-f) -#2, (g-i)-# 4, (j-l) -#5, (m-o)-# 6, (p-r)-# 7. The yellow dashed line represents the initial configuration of the wire.

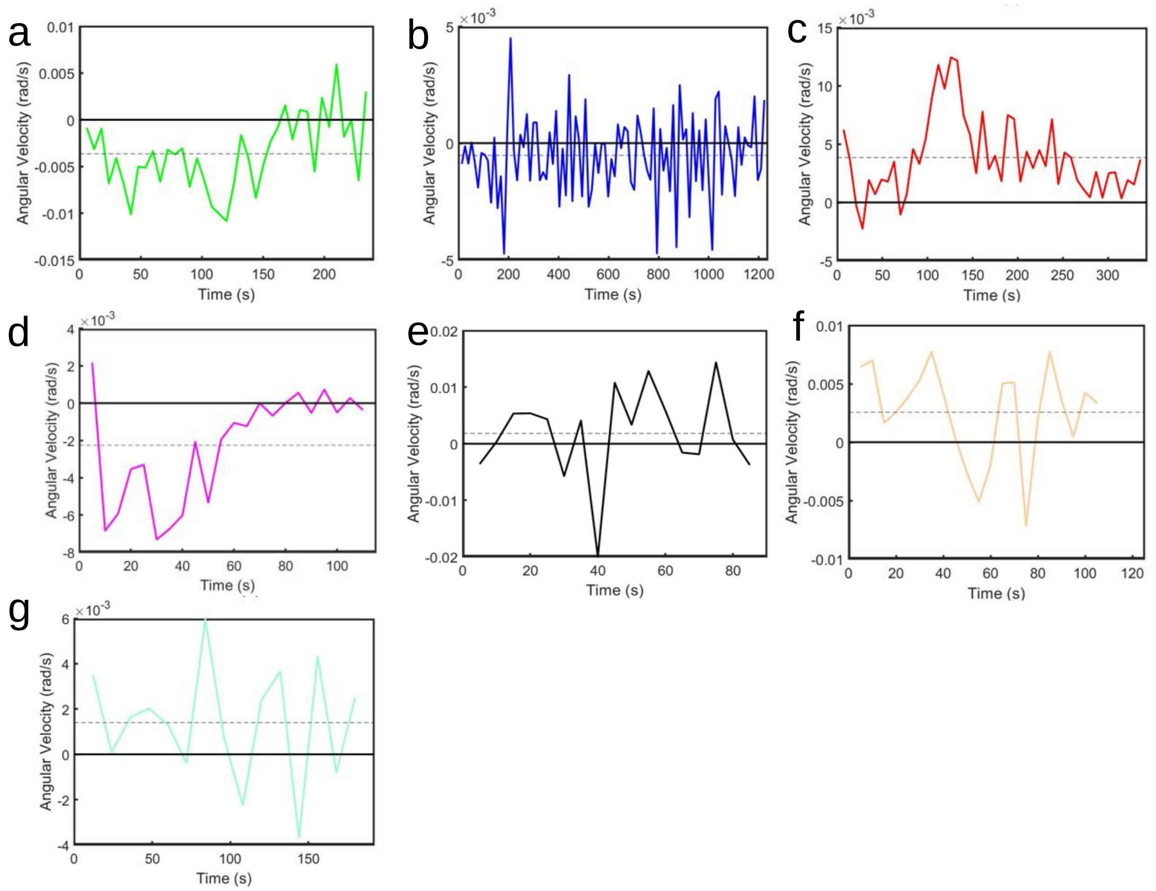

**Figure S2:** Angular velocity as a function of time for rotating nanowires, corresponding to the same color as listed in Table S1. The solid black line in each graph indicates zero value for the y-axis variable. The black dashed line indicates the average value of the y-axis variable across the time-series.

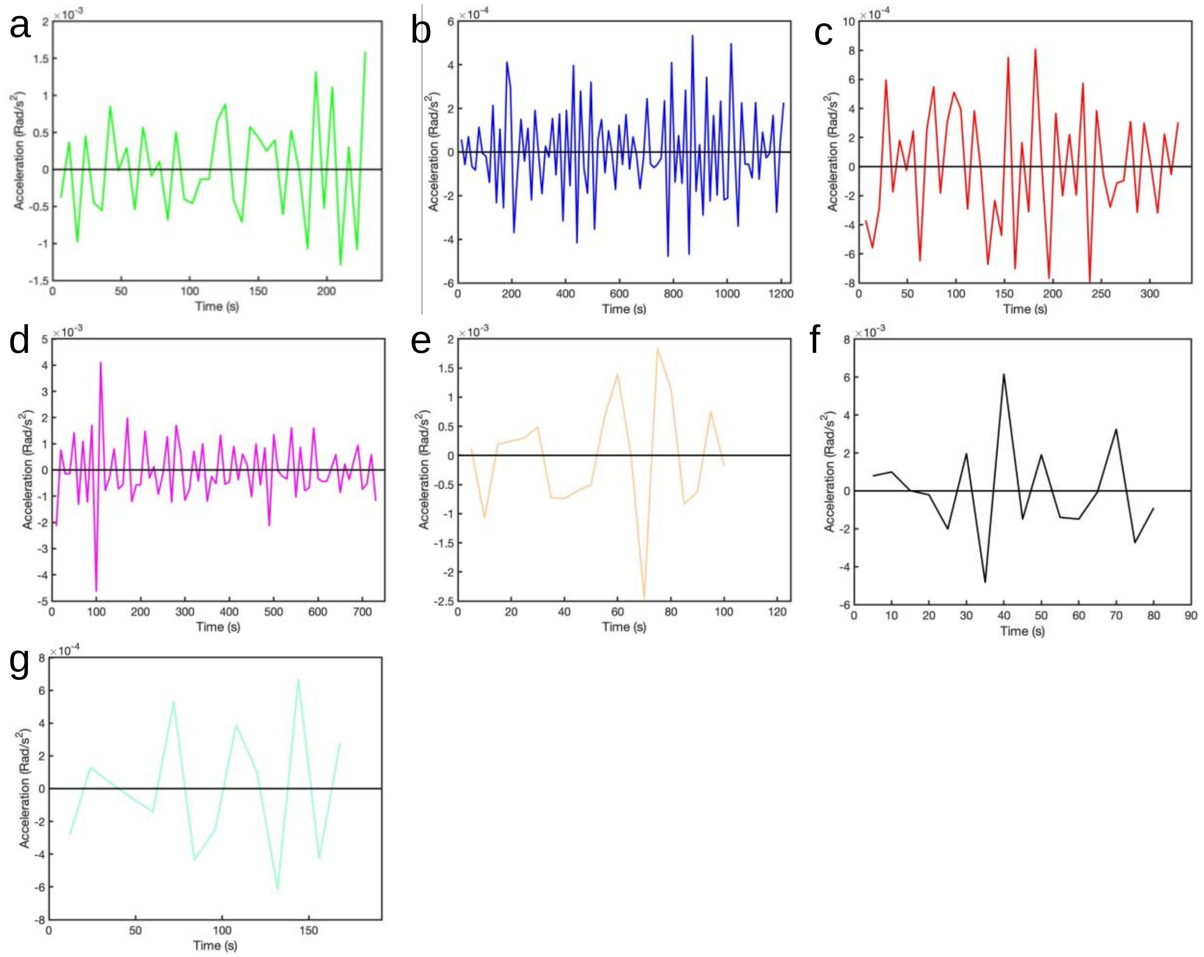

**Figure S3:** Angular acceleration as a function of time for rotating nanowires, corresponding to the same color as listed in Table S1. The solid black line in each graph indicates zero value for the y-axis variable.

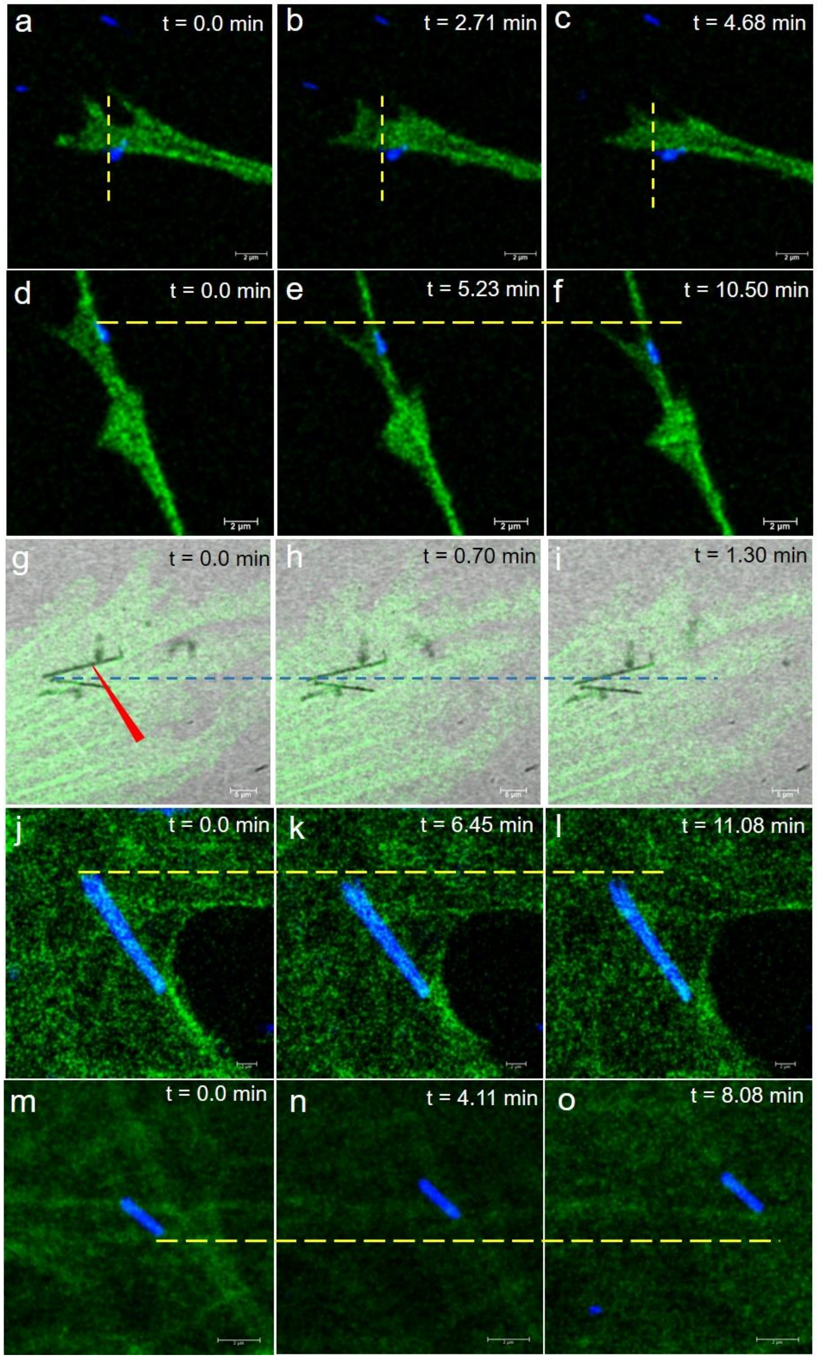

**Figure S4:** Snapshots of nanowire sliding along lamella or filopodia with coordinated assistance from membrane projections on those surfaces. Nanowires correspond to the # as listed in Table S2: (a-c) -#2,(d-f)-# 3, (g-i)-# 4, (j-l)-# 5, (m-o)-# 6. The yellow dashed line represents the initial configuration of the wire.

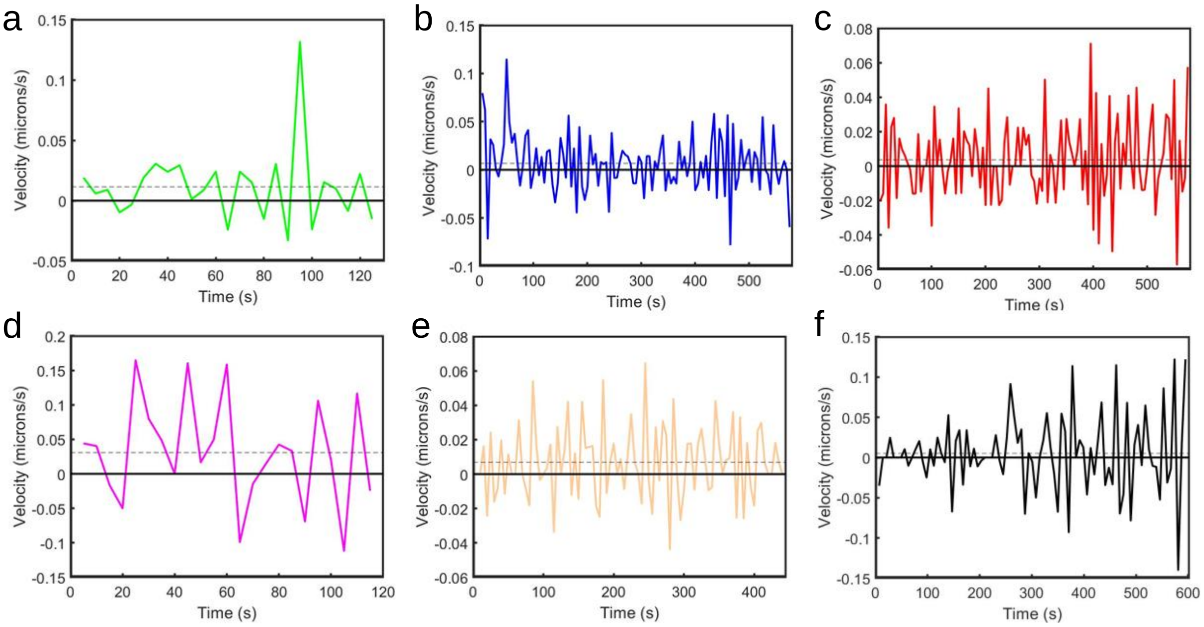

**Figure S5:** Velocity as a function of time for rotating nanowires, corresponding to the same color as listed in Table 2. The solid black line in each graph indicates zero value for the y-axis variable. The dashed black line represents the average value for the y-axis variable.

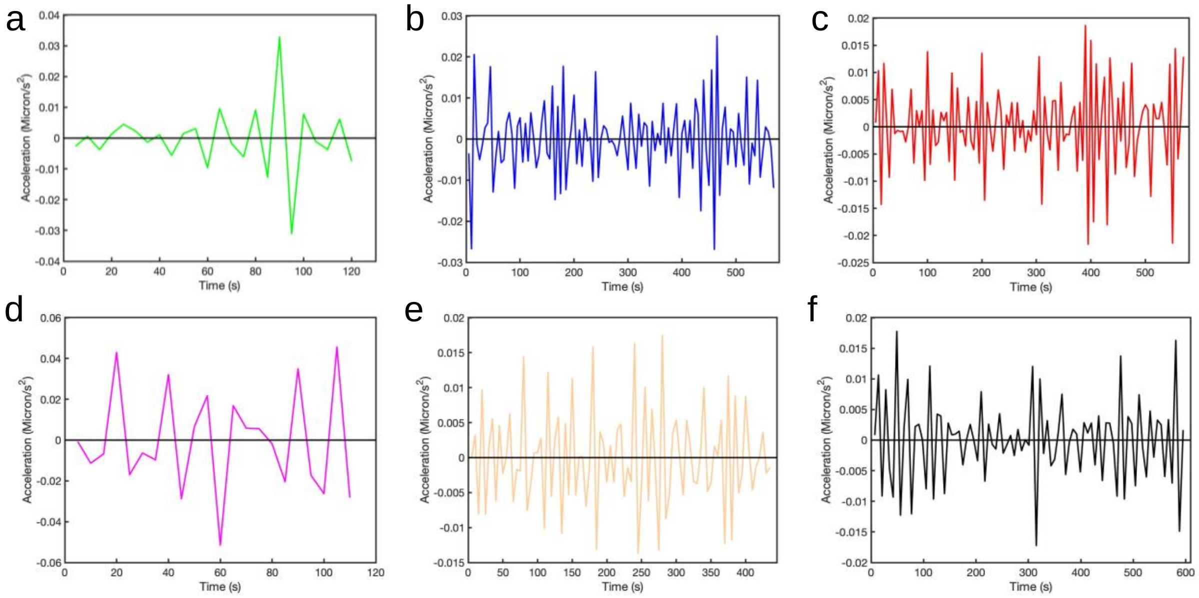

**Figure S6:** Acceleration as a function of time for rotating nanowires, corresponding to the same color as listed in Table S2. The solid black line in each graph indicates zero value for the y-axis variable.

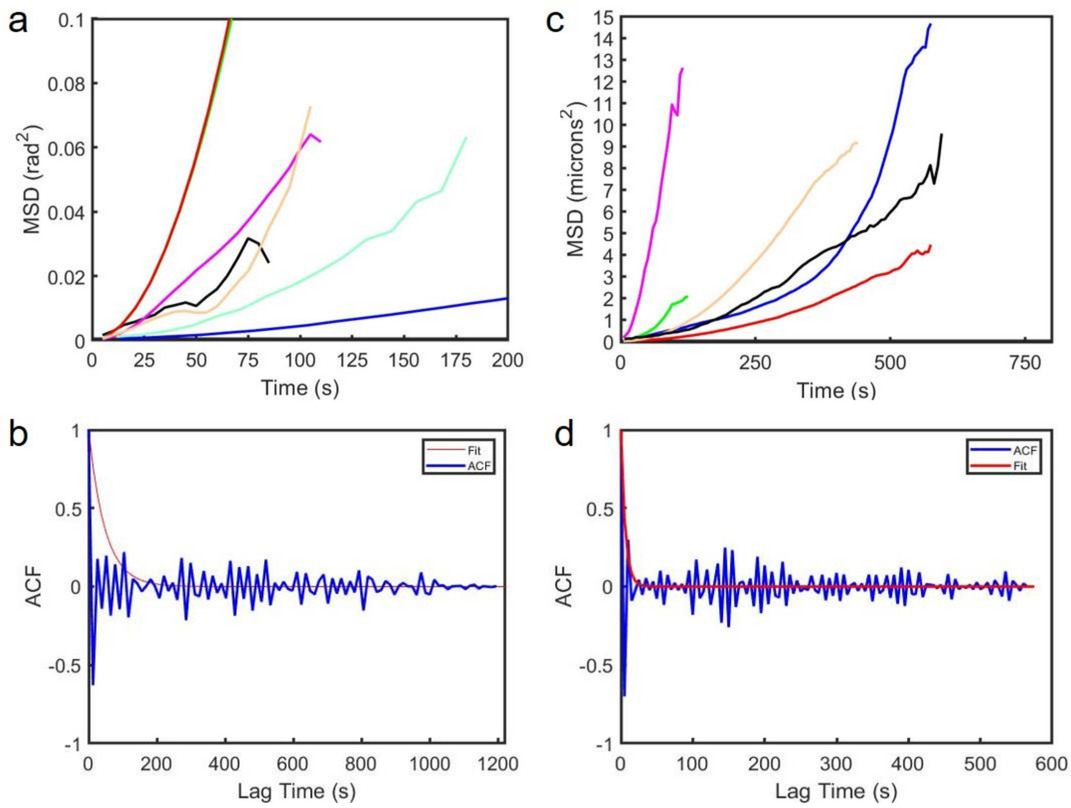

**Figure S7:** a. Rolling frame mean square displacement (MSD) of various nanowires tracked for their rotation to align with filopodia or rotation on a lamella. b. Representative autocorrelation function (ACF) of noise in angular velocity of a nanowire undergoing rotation to align with a filopodia or lamella. The exponential decay fit (Red) to the ACF is used to estimate the noise correlation time. c. Rolling mean square displacement (MSD) of various nanowires sliding along a filopodia or lamella. d. Representative autocorrelation function (ACF) of noise in velocity of a nanowire sliding along a filopodia or lamella. The exponential decay fit (Red) to the ACF is used to estimate the noise correlation time.

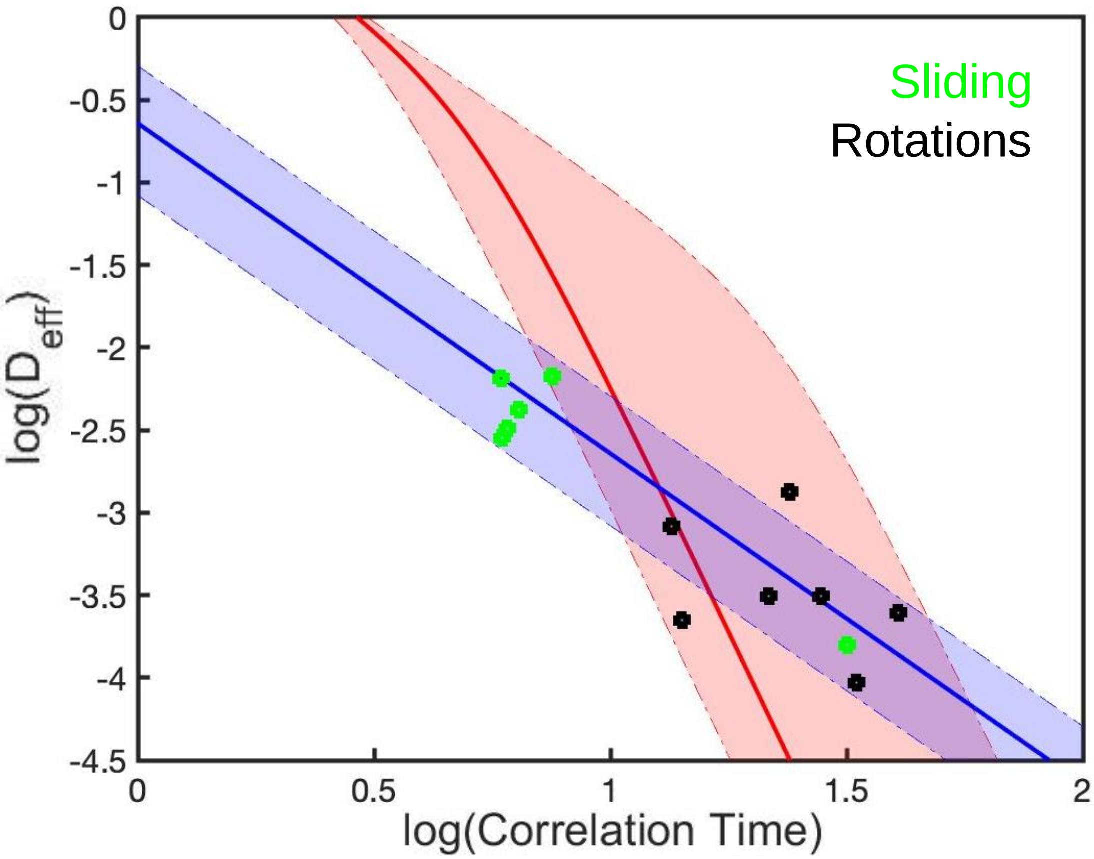

**Figure S8:** Comparison of the sliding wires versus rotating ones in the context of the OUP simulation.
